# Ascorbate maintains a low plasma low oxygen level

**DOI:** 10.1101/2020.02.06.937037

**Authors:** Louise Injarabian, Marc Scherlinger, Anne Devin, Stéphane Ransac, Jens Lykkesfeldt, Benoit S Marteyn

**Author notes:** Corresponding author: Benoit Marteyn, Institut de Biologie Moléculaire et Cellulaire Pasteur, 15, rue Descartes 67000 Strasbourg France.

## Abstract

In human blood, oxygen is mainly transported by red blood cells. Accordingly, the oxygen level in plasma is expected to be limited, although it has not been quantified yet. Here, by developing dedicated methods and tools, we determined that human plasma pO_2_ = 8.4 mmHg (1.2% O_2_). Oxygen solubility in plasma was believed to be similar to water. Here we reveal that plasma has an additional ascorbate-dependent oxygen-reduction activity. Plasma oxygenation oxidizes ascorbate (49.5 μM in fresh plasma vs <2 μM in oxidized plasma) and abolishes this capacity, which is restored by ascorbate supplementation. We confirmed these results *in vivo*, showing that the plasma pO_2_ is significantly higher in ascorbate-deficient guinea pigs (plasma ascorbate < 2μM), compared to control (plasma ascorbate > 15μM). Plasma low oxygen level preserves the integrity of oxidation-sensitive components such as ubiquinol. Circulating leucocytes are well adapted to these conditions, since the abundance of their mitochondrial network is limited.

These results shed a new light on the importance of oxygen exposure on leucocyte biological study, in regards with the reducing conditions they encounter *in vivo*; but also on the manipulation of blood products to improve their integrity and potentially improve transfusions’ efficacy.

**Key points:** 

## Introduction

Blood plasma is composed of water, ions, organic molecules, such as proteins, and vitamins. Gases, such as carbon dioxide (CO_2_) and oxygen (O_2_) also enter into the plasma composition, according to Henry’s law. The solubility coefficient of O_2_ (α_O2_) in plasma at 37°C is low compared to the CO_2_ (α_O2_ = 0.0031 mL O_2_/mmHg/100 mL blood vs α_CO2_ = 0.069 mL CO_2_/mmHg/100 mL blood)^1^. Only a limited fraction of O_2_ is dissolved in plasma, representing less than 2% of the total blood oxygen content. Arterial pO_2_ equals 75-100 mmHg and venous pO_2_ equals 30-50 mmHg; in theory the blood plasma pO_2_ would be ranged from 0.9 to 3 mmHg. The plasma fraction is anticipated to be poorly oxygenated in the overall blood circulation, although it has not been experimentally quantified. Until now, it was considered that the solubility coefficient of O_2_ in plasma was similar in water or saline^2^. The impact of ascorbate (or Vitamin C), a strong reducing molecule, on plasma oxygen level has not yet been investigated, despite of its abundance (plasma ascorbate concentration 50-70 μM^3,4^) and the respective standard redox potential of O_2_ and ascorbate (E’^0^_O2/H2O_ =0.815 and E’^0^_DHA/Ascorbate_ =0.08 at 25°C, P=1atm, pH=7)^5^.

In this report, by developing innovative strategies and tools we assessed plasma oxygen level and stability for the first time. We confirmed experimentally that plasma is poorly oxygenated and revealed that ascorbate contributes to its low oxygenation level, by reducing O_2_. The impact of plasma “physiological hypoxia” on circulating cells’ physiology and components’ stability and redox status has been further investigated.

## Methods

### Blood collection tubes

Blood samples were collected either in commercial collection tubes (BD Vacutainer K2E (EDTA), ref 368861) or in Hypoxytubes developed in collaboration with the Greiner Bio One (GBO) company, containing a limited amount of O_2_. Internal pO_2_ was quantified in commercial tubes and in Hypoxytubes using an oximeter with a microsensor equipped with a steel needle (Unisense).

### Blood collection

All participants gave written informed consent in accordance with the Declaration of Helsinki principles. Human blood was collected from healthy patients at the ICAReB service of the Pasteur Institut (authorization No. 2020_0120).

### Cell culture

HEK293T (ATCC CRL-1573) and Hep-G2 (ATCC HB-8065) were cultured in DMEM + 8% SVF. Cells were seeded onto 24-well plates and incubated 24h at 37°C at 0% (anoxic cabinet) or 21% O_2_.

White blood cells (WBCs) were purified form whole blood in an anoxic chamber by the addition of a 6% dextran solution (30 min, RT). The WBC-containing supernatant was collected and resuspended in RPMI 1640 (Thermofisher); remaining red blood cells were eliminated with a lysis buffer.

Cells were fixed in paraformaldehyde (PFA) 3.3% for immunofluorescent labelling or labeled with fluorescent marker for flow cytometry analysis, as previously described^6^.

### Plasma pO_2_ measurement and components’ dosage

Immediately after blood collection, the plasma pO_2_ was measured directly in the blood collection tube using an oximeter with a standardized microsensor equipped with a steel needle (Unisense), as previously described^7^.

Following centrifugation for 5 min at 2,000 *x g*, the plasma was acidified with an equal volume of 10% (w/v) metaphosphoric acid (MPA) containing 2 mmol/L of disodium-EDTA. Ascorbate concentration was quantified by high-performance liquid chromatography with coulometric detection, as described previously^8^. Likewise, using high-performance liquid chromatography with coulometric detection, α- and γ-tocopherol were analyzed as described by Sattler *et al*.^9^, and ubiquinone and ubiquinol as described elsewhere^10^.

Plasma potassium, calcium, magnesium, albumin, fibrinogen, Factor V and Factor VIII were quantified by a medical laboratory (Cerballiance, Paris, France).

### Plasma oxygen reduction rate quantification

Plasma oxygen consumption rate was measured with an oxymeter (Oroboros O2k-FluoRespirometer). Immediately after blood collection, samples were centrifuged, and plasma fractions were loaded in closed cuves (2 mL). Oxygen consumption fluxes were assessed when reaching constant values. Experiments were conducted with fresh plasma and after oxidation (exposure to atmospheric air: at least 30 min on a rotator mixer).

### Mitochondria study

#### Imaging

Mitochondria were immunolabeled with anti-SDHA antibody (ab14715, Abcam) in combination with a conjugated Alexa Fluor-568 (2124366, Invitrogen) on fixed preparations. Nuclei were labeled with DAPI. Cell imaging was performed with a confocal microscope (Leica DM5500 TCS SPE).

#### Flow cytometry

Cells were resuspended in PBS + 2 mM EDTA, labeled with 100 nM TMRM (T5428, Sigma-Aldrich) and analyzed with FACSCcalibur (BD Biosciences). Data were quantified with the FlowJo software (FlowJo, LLC).

### Guinea pig plasma analysis

3-week Dunkin-Hartley guinea pigs (Charles River) were fed for fifteen days with a standard diet (400 mg ascorbate/kg, Safediet ref. 106) or an ascorbate-deficient diet (<50 mg ascorbate/kg). Blood samples were collected in Hypoxytubes; plasma ascorbate concentration and pO_2_ were determined as described above. Procedure approved by the Institut Pasteur ethics committee (auth. n°190127).

### Statistics

Data were analyzed with the Prism 8 software (GraphPad). ANOVA or Student T-test were performed to analyze the different datasets.

## Results and Discussion

### Blood plasma is poorly oxygenated

Since all commercial tubes contain a significant amount of oxygen (here, 75.7 ± 4.6 mmHg), we designed and produced tubes containing a limited amount of oxygen (15.9 ± 2.9 mmHg) hereafter termed Hypoxytube (Figure 1A), to avoid experimental oxygenation of blood samples. Oxygen level in plasma was quantified with a needle sensor in commercial tubes or Hypoxytubes (Figure 1B). Plasma pO_2_ was 9.8 ± 4.8 mmHg in commercial tubes versus 8.4 ± 1.0 mmHg in Hypoxytube (*p*<0.01): the lastest value being the most accurate quantification of the plasma oxygen level so far (Figure 1C).

**Figure 1.**
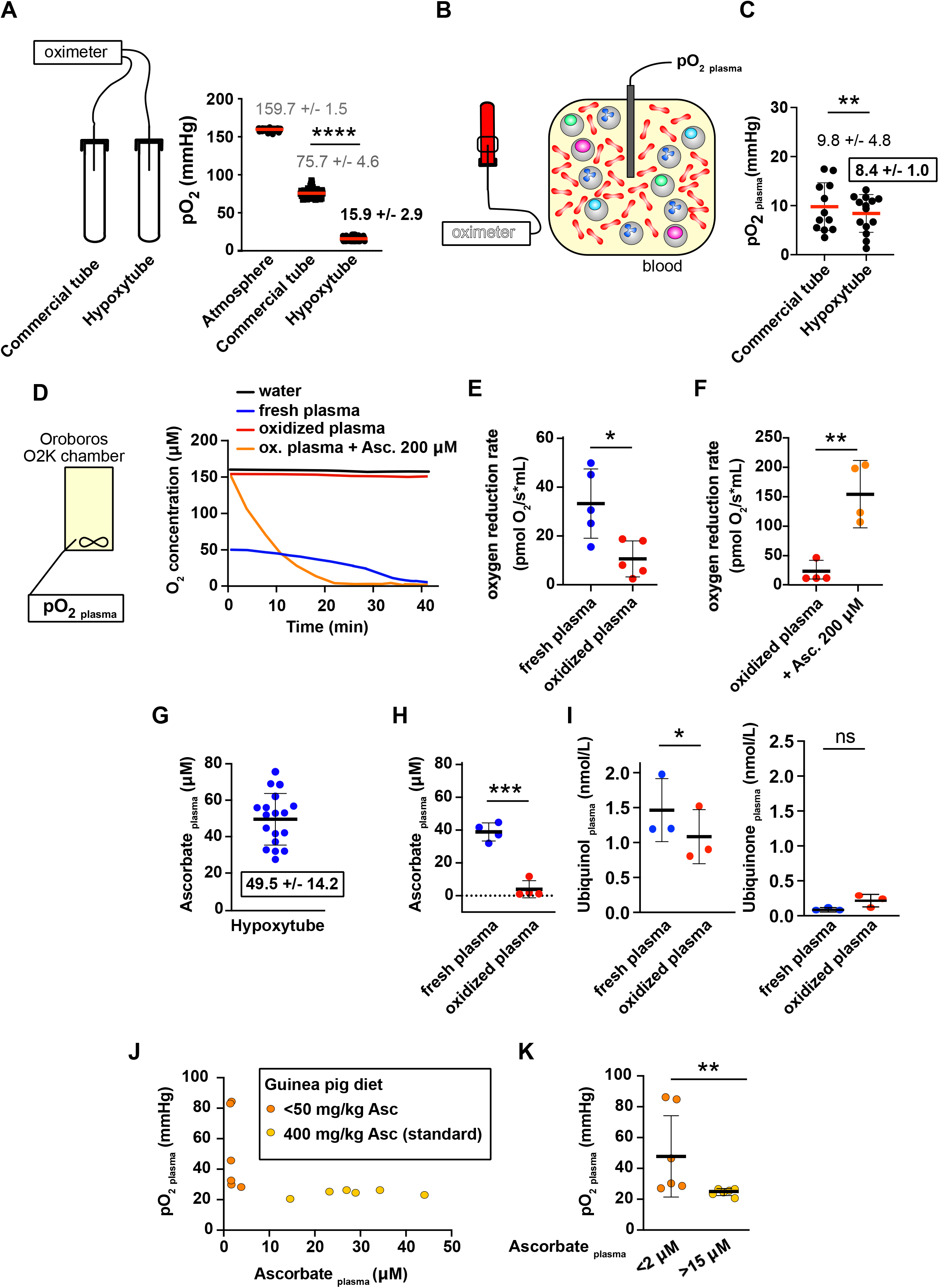
The plasma oxygen level is low, mainly sustained by the ascorbate oxygen. **(A)** Blood collection tubes containing a limited amount of oxygen (Hypoxytubes) have been designed and validated using an oximeter with a microsensor equipped with a steel needle. Commercial tube used as a control was BD Vacutainer K2E (EDTA). Results are expressed as Mean ± S.D.; **** indicates p<0.0001, n=20. **(B-C)** Plasma oxygen level was directly quantified in whole venous blood collected in commercial tube or Hypoxytube. Plasma pO_2_ quantifications are expressed as Mean ± S.D.; ** indicates p<0.01, n=12. **(D)** Plasma samples were loaded in closed cuve to record the time-dependent oxygen availability in fresh plasma, oxidized plasma supplemented or not with 200 μM ascorbate, and water (representative experiment). **(E)** Plasma oxygen reduction rates were quantified in fresh or oxidized plasma samples, as described in (D). Results are expressed as Mean ± S.D.; * indicates p<0.05, n=5. **(F)** The impact of oxidized plasma ascorbate supplementation (200 μM) was quantified as described in (E). Results are expressed as Mean ± S.D.; ** indicates p<0.01, n=4. **(G-H)** Plasma ascorbate concentration in fresh samples was quantified as described in Methods, in blood samples collected in Hypoxytubes (G, n=18). The impact of plasma oxygenation on ascorbate concentration is shown in (H, n=4). Results are expressed as Mean ± S.D.; *** indicates p<0.001. **(I)** Ubiquinol concentration in fresh and oxidized plasma samples was quantified, together with other plasma components (see Figure S1). Results are expressed as Mean ± S.D.; * indicates p<0.05, n=3. **(J-K)** The impact of plasma ascorbate deficiency on the control of the oxygen level has been investigated in vivo in guinea pigs (J-K). Plasma ascorbate concentration and pO_2_ was recorded in animals fed standard (400 mg ascorbate/kg) or ascorbate-deficient diet (<50 mg ascorbate/kg) (J). Plasma pO_2_ was average in each group (plasma ascorbate < 2μM (deficient) and > 15μM (control)). Results are expressed as Mean ± S.D.; ** indicates p<0.01, n=6.

### Ascorbate sustains a plasma oxygen-reduction activity

Oxygen solubility has so far been believed to be similar in plasma and water. However, when fresh plasma pO_2_ was recorded in a closed chamber (Oroboros), a continuous decrease was observed until anoxia was reached (Figure 1D). This reaction was significantly lower in oxidized plasma (exposed to atmospheric oxygen) or in water (Figure 1D-E). These results strongly suggested that an oxygen-sensitive plasma component was mediating its oxygen-reduction capacity. We hypothesized that plasma ascorbate may play a central role in this reaction. The supplementation of oxidized plasma with 200 μM ascorbate restored its oxygen reduction activity (Figure 1D and 1F), supporting this hypothesis. This ascorbate-dependent reaction does not occur in water (Figure S1A). The plasma ascorbate concentration was first determined in a large number of individuals (49.5 ± 14.2 μM, Figure 1G). We further demonstrated that plasma ascorbate concentration was drastically reduced in oxidized plasma compared to fresh plasma (*p*<0.001, Figure 1H), confirming ascorbate susceptibility to oxidation as previously reported. We further demonstrated that the concentration of ubiquinol, another oxidation-sensitive plasma component was significantly lower in oxidized plasma compared to fresh plasma (*p*<0.05, Figure 1I); this reaction was associated with an increase of ubiquinone (ubiquinol oxidized form), as expected (Figure 1I). The concentration of other plasma components was not modified by plasma oxygenation, including salts (potassium, calcium, magnesium, Figure S1B),proteins (albumin, fibrinogen, coagulation Factor V and VIII, Figure S1C) or additional oxidation-sensitive components (α-tocopherol, γ-tocopherol) (Figure S1D).

We confirmed the ascorbate-dependent plasma oxygen reduction capacity *in vivo*, in a guinea pig model. Like humans, guinea pigs do not synthesize ascorbate, and are consequently dependent on dietary supply. When animals were fed a standard diet (400 mg/kg ascorbate), the plasma ascorbate concentration was higher than 15 μM and the plasma pO_2_ controlled at a low level (24.11 ± 2.23 mmHg, Figure 1J-K). These values are comparable to human plasma although higher, probably due to technical reasons (increased lag time between blood collection and pO_2_ measurement). When animals were fed an ascorbate-deficient diet (<50 mg ascorbate/kg), the plasma ascorbate concentration was lower than 2 μM and the plasma pO_2_ no longer maintained at a low level (50.40 ± 26.32 mmHg) (Figure 1J-K). Altogether these results confirm the *in vivo* contribution of ascorbate to the maintenance of a low plasma oxygen level.

### Circulating leucocytes sense plasma low-oxygenation - mitochondrial network

The adaptation of circulating leucocytes to plasma low oxygen level has not previously been investigated. In other cell-types, it has been reported that under hypoxic conditions, mitochondrial abundance and oxygen consumption is reduced ^11–13^. In addition, mitochondria play a key function in leucocyte metabolism and function, beyond metabolism regulation^14^. We confirmed by immunofluorescence (Figure 2A) and flow cytometry (Figure 2B-D) that, compared to two different cell lines (HEK293T and HEp-2) cultured under atmospheric conditions, the mitochondrial abundance of leucocytes (granulocytes, monocytes and lymphocytes) was significantly reduced (ANOVA, Figure 2C-D). Cell lines’ mitochondrial content was significantly reduced upon culture under anoxic conditions (Figure S2C). These results confirm that leucocytes evolve under low oxygen conditions in the blood plasma fraction.

**Figure 2.**
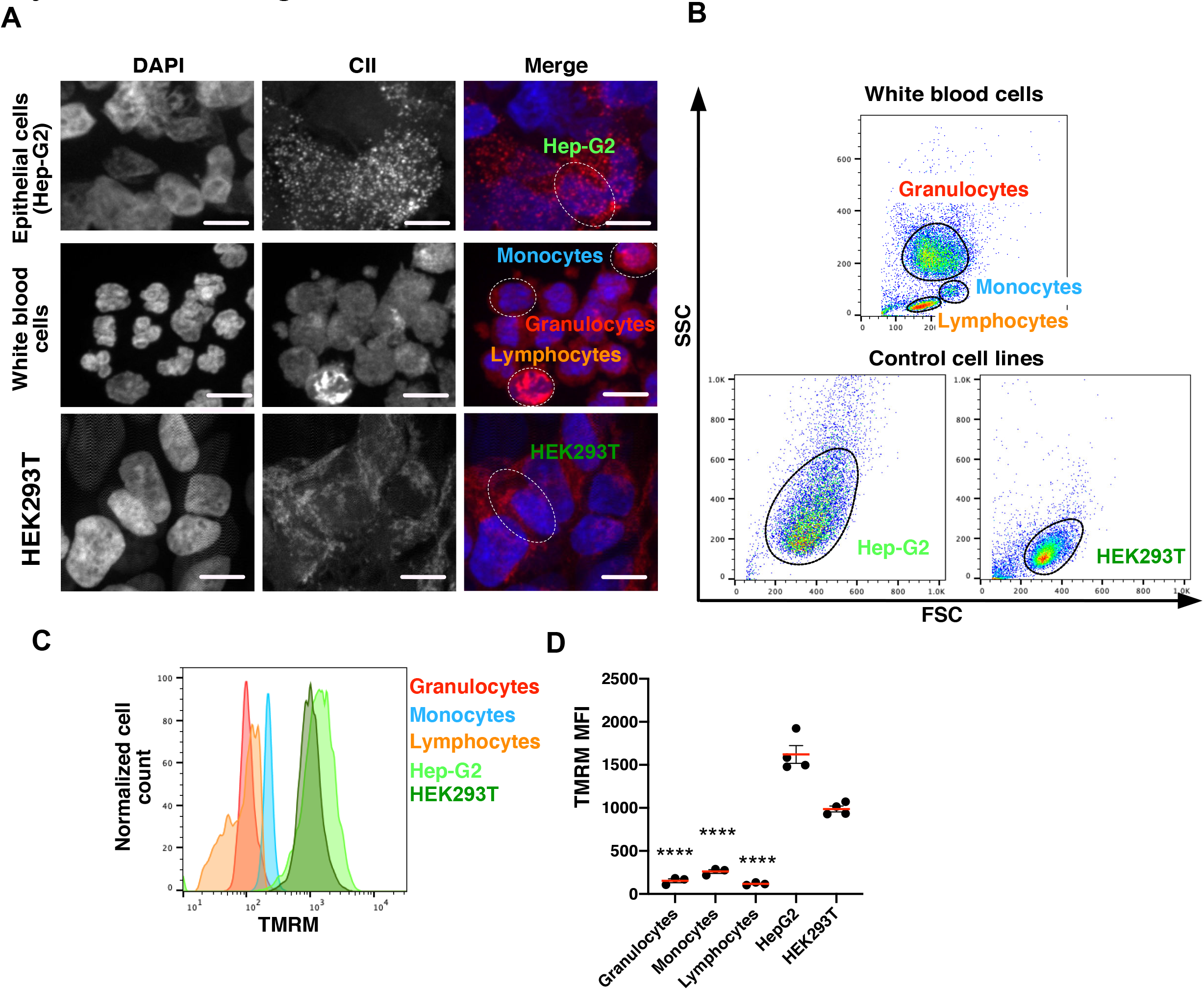
The mitochondria abundance is reduced in circulating leucocytes in low-oxygenated plasma. **(A)** Immunofluorescence staining of white blood cells (WBCs: monocytes, lymphocytes and granulocytes) Hep-G2 cells, and HEK293T cells using anti-CII (mitochondria, red) and DAPI (nuclei, blue). Bars are 10 μm. **(B)** Flow cytometry analysis of WBCs (granulocytes, monocytes and lymphocytes), Hep-G2 and HEK293T. Representation of SSC and FSC profiles. **(C)** TMRM (mitochondria) intensity profiles in granulocytes, monocytes, lymphocytes, Hep-G2 and HEK293T cells (representative experiment) **(D)** Quantification of TMRM mean fluorescence intensity (MFI) in cells described in (C). Results are expressed as Mean ± S.D.; **** indicates p<0.0001 (One-way ANOVA with Tukey’s test, see Tables in Figure S1A), n=4. See additional quantification in Figure S2C.

### Blood plasma low oxygenation level should be further considered for basic research and therapeutic application

Blood plasma low oxygen level should be considered more carefully to study blood cells in vitro.

It may have some important drawback in blood product collection and preservation prior transfusion to avoid detrimental impact on its quality.

Further investigations will be required to determine the intermediate reactions involved in the reduction of O_2_ by ascorbate in the plasma, potentially involving the transient formation of reactive oxygen species or ascorbate free radical^5^.

Plasma transfusion, preservation of samples from oxidation: red blood cells ^15^, platelet^16^

## Acknowledgements

This work was supported by the French National Research Agency: ANR JCJC 2017-17-CE15-0012 (BSM).

## Authorship contributions

LI conducted quantitative analysis of the data. MS performed experiments. JL quantified plasma ascorbate in plasma samples. LI, AD and SR contributed to data interpretation. BSM designed and performed the experiments, interpreted the data, and wrote the manuscript. All authors critically edited the draft manuscript and approved the final version.

## Disclosure of Conflicts of Interest

Authors declare no conflict of interest

**Supplementary data. Figure 2.**

**(A)** Water oxygen reduction rate w/wo Asc. **(B)** Plasma component quantification in fresh an oxidized plasma samples

**Supplementary data. Figure 2.**

**(A)** Results of ANOVA statistical analysis (Tukey’s multiple comparisons test) of TMRM MFI between granulocytes, monocytes, lymphocytes, HepG2 and HEK293T cells (Figure 2D), representing mean difference, 95% confidence interval of the difference, significance, summary and adjusted p-value. **(B)** Details of ANOVA statistical analysis test. **(C)** TMRM MFI in HepG2 and HEK293T cells exposed to atmospheric oxygen (21% O2) or anoxia (0% O2) for 24h at 37°C measured with flow cytometry. **(D)** Representation of TMRM profiles of HepG2 and HEK293T cells exposed or not to atmospheric oxygen (21%).

## References

1. Pittman RN. Regulation of Tissue Oxygenation. Colloquium Ser Integr Syst Physiology Mol Funct. 2011;3(3):1–100.

2. Christmas KM, Bassingthwaighte JB. Equations hfor O2 and CO2 Solubilities in Saline and Plasma: Combining Temperature and Density Dependences. J Appl Physiology Bethesda Md 1985. 2017;122(5):jap.01124.2016.

3. VanderJagt D, Garry P, Hunt W. Ascorbate in plasma as measured by liquid chromatography and by dichlorophenolindophenol colorimetry. Clin Chem. 1986;32(6):1004–6.

4. Lykkesfeldt J, Tveden-Nyborg P. The Pharmacokinetics of Vitamin C. Nutrients. 2019;11(10):2412.

5. Cabelli DE, Bielski BH. Kinetics and mechanism for the oxidation of ascorbic acid/ascorbate by HO2/O2- (hydroperoxyl/superoxide) radicals. A pulse radiolysis and stopped-flow photolysis study. J Phys Chem. 1983;87(10):1809–1812.

6. Monceaux V, Chiche-Lapierre C, Chaput C, et al. Anoxia and glucose supplementation preserve neutrophil viability and function. Blood. 2016;128(7):993–1002.

7. Tinevez J-Y, Arena ET, Anderson M, et al. Shigella-mediated oxygen depletion is essential for intestinal mucosa colonization. Nat Microbiol. 2019;4(11):2001–2009.

8. Lykkesfeldt J. Measurement of ascorbic acid and dehydroascorbic acid in biological samples. Curr Protoc Toxicol Éditor Board Mahin D Maines Ed Et Al. 2002;Chapter 7(1):Unit 7.6.1–15.

9. Sattler W, Mohr D, Stocker R. Rapid isolation of lipoproteins and assessment of their peroxidation by high-performance liquid chromatography postcolumn chemiluminescence. Methods Enzymol. 1994;233:469–89.

10. Schou-Pedersen AV, Schemeth D, Lykkesfeldt J. Determination of Reduced and Oxidized Coenzyme Q10 in Canine Plasma and Heart Tissue by HPLC-ECD: Comparison with LC-MS/MS Quantification. Antioxidants. 2019;8(8):253.

11. Fukuda R, Zhang H, Kim J, et al. HIF-1 Regulates Cytochrome Oxidase Subunits to Optimize Efficiency of Respiration in Hypoxic Cells. Cell. 2007;129(1):111–122.

12. Zhang H, Gao P, Fukuda R, et al. HIF-1 Inhibits Mitochondrial Biogenesis and Cellular Respiration in VHL-Deficient Renal Cell Carcinoma by Repression of C-MYC Activity. Cancer Cell. 2007;11(5):407–420.

13. Papandreou I, Cairns RA, Fontana L, Lim A, Denko NC. HIF-1 mediates adaptation to hypoxia by actively downregulating mitochondrial oxygen consumption. Cell Metab. 2006;3(3):187–197.

14. Mehta MM, Weinberg SE, Chandel NS. Mitochondrial control of immunity: beyond ATP. Nat Rev Immunol. 2017;17(10):608–620.

15. Mohanty JG, Nagababu E, Rifkind JM. Red blood cell oxidative stress impairs oxygen delivery and induces red blood cell aging. Front Physiol. 2014;5:84.

16. Manasa K, Vani R. Influence of Oxidative Stress on Stored Platelets. Adv Hematology. 2016;2016:4091461.

