## Supplementary figures and images for "Ascorbate maintains a low plasma low oxygen level"

### Supplementary Figure 1

**Injarabian L et al, Supplementary Figure 1**

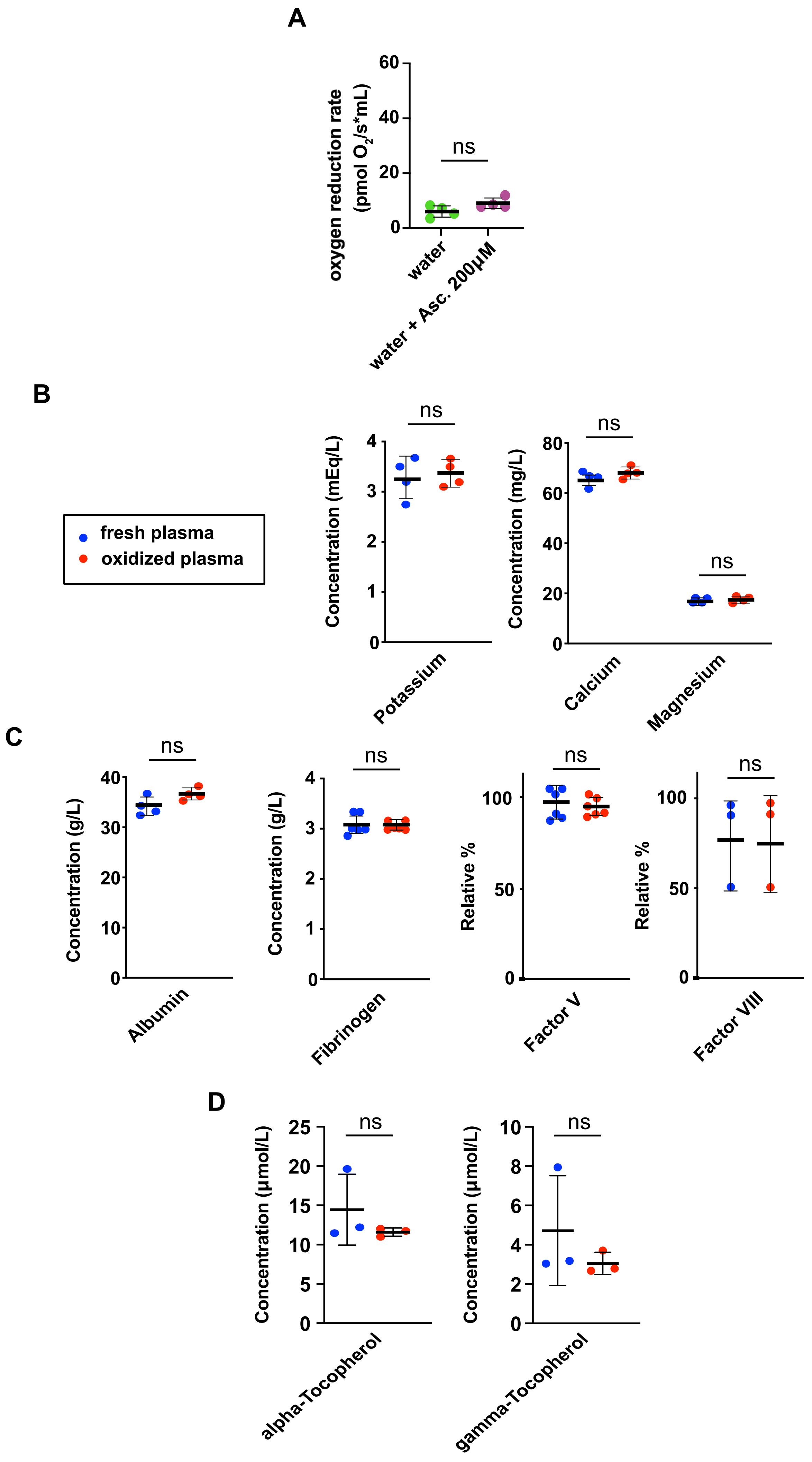
