## Supplementary Figure 2 for "Ascorbate maintains a low plasma low oxygen level"

**Injarabian L et al, Supplementary Figure 2**

**A**

| Tukey's multiple comparisons test | Mean Diff. | 95.00% CI of diff. | Significant? | Summary | Adjusted P value |
|---|---|---|---|---|---|
| Granulocytes vs. Monocytes | -107,3 | -397.5 to 182.8 | No | ns | 0,7627 |
| Granulocytes vs. Lymphocytes | 35,33 | -254.8 to 325.5 | No | ns | 0,9945 |
| Granulocytes vs. Hep-G2 | -1468 | -1739 to -1197 | Yes | **** | <0.0001 |
| Granulocytes vs. HEK293T | -834,2 | -1106 to -562.8 | Yes | **** | <0.0001 |
| Monocytes vs. Lymphocytes | 142,7 | -147.5 to 432.8 | No | ns | 0,5426 |
| Monocytes vs. Hep-G2 | -1361 | -1632 to -1089 | Yes | **** | <0.0001 |
| Monocytes vs. HEK293T | -726,8 | -998.2 to -455.5 | Yes | **** | <0.0001 |
| Lymphocytes vs. Hep-G2 | -1503 | -1775 to -1232 | Yes | **** | <0.0001 |
| Lymphocytes vs. HEK293T | -869,5 | -1141 to -598.1 | Yes | **** | <0.0001 |
| Hep-G2 vs. HEK293T | 633,8 | 382.5 to 885.0 | Yes | **** | <0.0001 |

**B**

| Test details | Mean 1 | Mean 2 | Mean Diff. | SE of diff. | n1 | n2 | | DF |
|---|---|---|---|---|---|---|---|---|
| Granulocytes vs. Monocytes | 153,3 | q 260,7 | -107,3 | 91,02 | 3 | 3 | 1,67 | 12 |
| Granulocytes vs. Lymphocytes | 153,3 | 118 | 35,33 | 91,02 | 3 | 3 | 0,55 | 12 |
| Granulocytes vs. Hep-G2 | 153,3 | 1621 | -1468 | 85,14 | 3 | 4 | 24,4 | 12 |
| Granulocytes vs. HEK293T | 153,3 | 987,5 | -834,2 | 85,14 | 3 | 4 | 13,9 | 12 |
| Monocytes vs. Lymphocytes | 260,7 | 118 | 142,7 | 91,02 | 3 | 3 | 2,22 | 12 |
| Monocytes vs. Hep-G2 | 260,7 | 1621 | -1361 | 85,14 | 3 | 4 | 22,6 | 12 |
| Monocytes vs. HEK293T | 260,7 | 987,5 | -726,8 | 85,14 | 3 | 4 | 12,1 | 12 |
| Lymphocytes vs. Hep-G2 | 118 | 1621 | -1503 | 85,14 | 3 | 4 | 25 | 12 |
| Lymphocytes vs. HEK293T | 118 | 987,5 | -869,5 | 85,14 | 3 | 4 | 14,4 | 12 |
| HepG2 vs. HEK293T | 1621 | 987,5 | 633,8 | 78,82 | 4 | 4 | 11,4 | 12 |

**C**

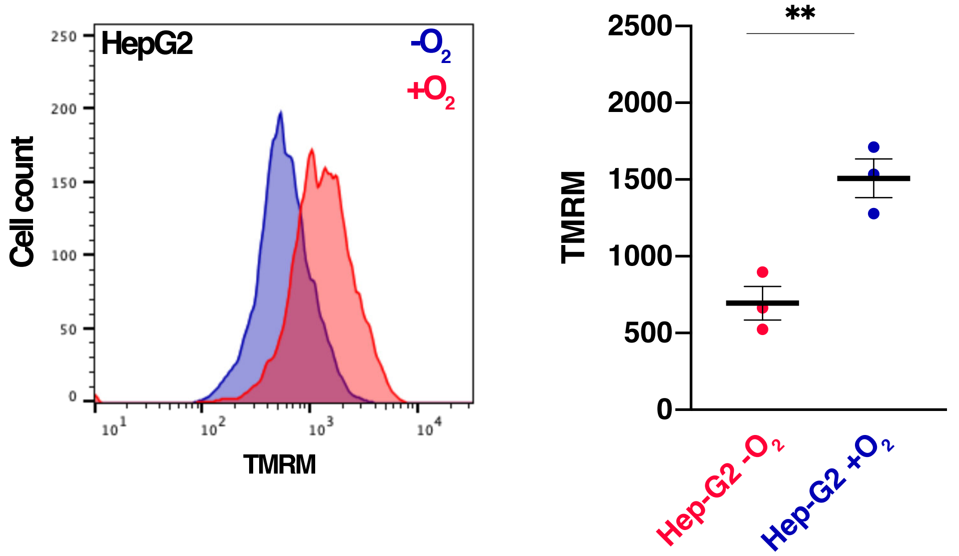
